# Analysis of 19 Heliconiine Butterflies Shows Rapid TE-based Diversification and Multiple SINE Births and Deaths

**DOI:** 10.1101/596502

**Authors:** David A Ray, Jenna R Grimshaw, Michaela K Halsey, Jennifer M Korstian, Austin B Osmanski, Kevin AM Sullivan, Kristen A Wolf, Harsith Reddy, Nicole Foley, Richard D Stevens, Binyamin Knisbacher, Orr Levy, Brian Counterman, Nathan B Edelman, James Mallet

**Affiliations:** Department of Biological Science, Texas Tech University, Lubbock, TX; Department of Veterinary Integrative Biosciences, College of Veterinary Medicine, Texas A&M University, College Station, TX; Department of Natural Resources Management, Texas Tech University, Lubbock, TX; The Mina and Everard Goodman Faculty of Life Sciences, Bar-Ilan University, Ramat Gan 52900, Israel; Broad Institute of MIT and Harvard, Cambridge, MA; Department of Physics, Bar-Ilan University, Ramat Gan 52900, Israel; Department of Biological Sciences, Mississippi State University, Mississippi State, MS; Department of Organismic and Evolutionary Biology, Harvard University, Cambridge, MA

## Abstract

Transposable elements (TEs) play major roles in the evolution of genome structure and function. However, because of their repetitive nature, they are difficult to annotate and discovering the specific roles they may play in a lineage can be a daunting task. Heliconiine butterflies are models for the study of multiple evolutionary processes including phenotype evolution and hybridization. We attempted to determine how TEs may play a role in the diversification of genomes within this clade by performing a detailed examination of TE content and accumulation in 19 species whose genomes were recently sequenced. We found that TE content has diverged substantially and rapidly in the time since several subclades shared a common ancestor with each lineage harboring a unique TE repertoire. Several novel SINE lineages have been established that are restricted to a subset of species. Furthermore, the previously described SINE, Metulj, appears to have gone extinct in two subclades while expanding to significant numbers in others. Finally, a burst of TE origination corresponds temporally to a burst of speciation in the clade, potentially providing support to hypotheses that TEs are drivers of genotypic and phenotypic diversification. This diversity in TE content and activity has the potential to impact how heliconiine butterflies continue to evolve and diverge.

## Introduction

TEs have been described as ‘drivers of genome evolution’ (Kazazian 2004). Indeed, transposable elements (TEs) are major contributors to processes that influence genomic change (Kidwell and Lisch 1997). TEs mediate small-scale changes but also influence large-scale structural changes including deletions, translocations, duplications, ectopic recombination and are intimately associated with the evolution of genome size in some lineages (Lim and Simmons 1994; Gray 2000; Hedges and Deininger 2007; Carbone, et al. 2014; Grabundzija, et al. 2016; Kapusta, et al. 2017). Transposition is an efficient mechanism for generating widespread genetic diversity that evolutionary processes may build on, leading to phenotypic and taxonomic diversity. While structural changes induced by the insertion of hundreds or thousands of 200 to 10,000 bp units at a time are likely important to evolutionary processes, it has also been argued that by contributing multiple copies of ready-to-use regulatory motifs, transposons also induce more subtle but also more significant (in the long run) regulatory innovation (Rebollo, et al. 2010; Rebollo, et al. 2012; Ellison and Bachtrog 2013; Jacques, et al. 2013; Sundaram, et al. 2014; Chuong, et al. 2016; Mita and Boeke 2016; Chuong, et al. 2017; Sundaram, et al. 2017; Trizzino, et al. 2017).

The idea that by generating genomic diversity TEs play a significant role in adaptive change is not new. On the contrary, the discoverer of TEs herself, Barbara McClintock, proposed that TEs may act as a mechanism for the genome to respond to stress in an adaptive manner (McClintock 1956, 1984). More recently, Oliver and Greene (2011, 2012) proposed the TE Thrust Hypothesis, suggesting that TEs enhance evolutionary potential by introducing variation in the genomes they occupy. In a related hypothesis, Zeh et al. (2009) suggested reduced epigenetic suppression of TEs when organisms are under stress, thereby increasing their activity and their impact on genome structure. This is referred to as the Epi-Transposon Hypothesis. Other authors have offered similar and/or related ideas, in every case linking transposon activity to adaptation (Jurka, et al. 2011; Jurka, et al. 2012; Koonin 2016a, b).

While these ideas represent significant advances in our understanding of TE-genome interactions, several limitations have restricted the scope of research on the relationship between TEs and diversification, preventing tests of these major hypotheses and generalization across taxa. First, the comparisons undertaken thus far involve relatively deep divergences that make understanding the changes that occur at lower taxonomic levels difficult to tease apart. Second, cost effective approaches to densely sample divergent clades have only become available recently, limiting prior comparisons to such deep divergences in an effort to maximize observable differences. Third, a mechanistic understanding of TE action has been confined to lab models and their cell lines, limiting research into the emergence and control of phenotypic traits. However, recent advances have created opportunities to move past these barriers. Primarily, cost reductions and advances in sequencing technologies and genome analysis have allowed us to examine larger and larger numbers of whole genomes, including whole genome comparisons among relatively closely related species (Lamichhaney, et al. 2015; Nater, et al. 2015). A narrowed focus has the potential to inform the scientific community of the influences TEs may have at the early stages of taxonomic divergence.

Butterflies of the genus *Heliconius* and related genera are models for the study of several evolutionary processes from hybridization to the evolution of Müllerian mimicry (Heliconius 2012). They have experienced multiple recent bursts of speciation and represent an adaptive radiation that is ripe for study at the genome level (Supple, et al. 2013; Supple, et al. 2014; Kozak, et al. 2015; Arias, et al. 2017). These characteristics create an excellent opportunity to examine patterns of TE evolution in a rapidly diversifying clade, allowing us to ask questions about how the TEs themselves evolve as species diverge from one another.

TEs from *Heliconius* were first described as part of the first *Heliconius melpomene* genome project (Heliconius 2012) and examined in detail by Lavoie et al. (2013). In that work, the TE landscape was revealed to be exceptionally diverse with large numbers of active LINEs (Long INterspersed Elements) and large genome proportions derived from SINEs (Short INterspersed Elements) and rolling circle transposons (Helitrons). Further, the genome was shown to be labile, especially with regard to larger TEs, which appear to be removed regularly via non-homologous recombination. This is in line with recent hypotheses related to genome evolution and TE content, and in particular, the accordion model of genome evolution, in which some DNA is contributed while other DNA is jettisoned over evolutionary time (Kapusta, et al. 2017).

Recently, multiple representatives from this clade were subjected to whole genome sequence analysis using multiple assembly technologies, providing us with an opportunity to examine the evolution of TEs in detail across 20 heliconiine species (Edelman et al. submitted). We performed de novo TE annotations on 19 of these genomes and compared the TE landscapes across the heliconiine tree, revealing patterns of transposable element evolution not yet seen at this fine a scale. We see that differential TE accumulation can be established rapidly across lineages and that particular families and subfamilies establish themselves differentially in independent lineages in relatively short periods of time.

This detailed examination of TE evolution in closely related species lays the groundwork for additional analysis of transposable elements as members of genomic communities that evolve in ways similar to natural communities in ecosystems. It also opens the door to examining genomic factors that may influence the relative success of TEs in each genome as they diverge from one another.

## Results

### Data

Draft genomes for 19 species were analyzed for transposable element content (Figure 1). Details of each assembly are available in Edelman et al. (submitted) and in Supplemental Table 1. One species, *H. melpomene*, has been analyzed thoroughly for TEs and therefore served as a starting point for some downstream analyses (Heliconius 2012; Lavoie, et al. 2013). We assumed that any old, shared insertions among the species analyzed were identified as part of that analysis or are part of other insect TE libraries.

**Figure 1.**
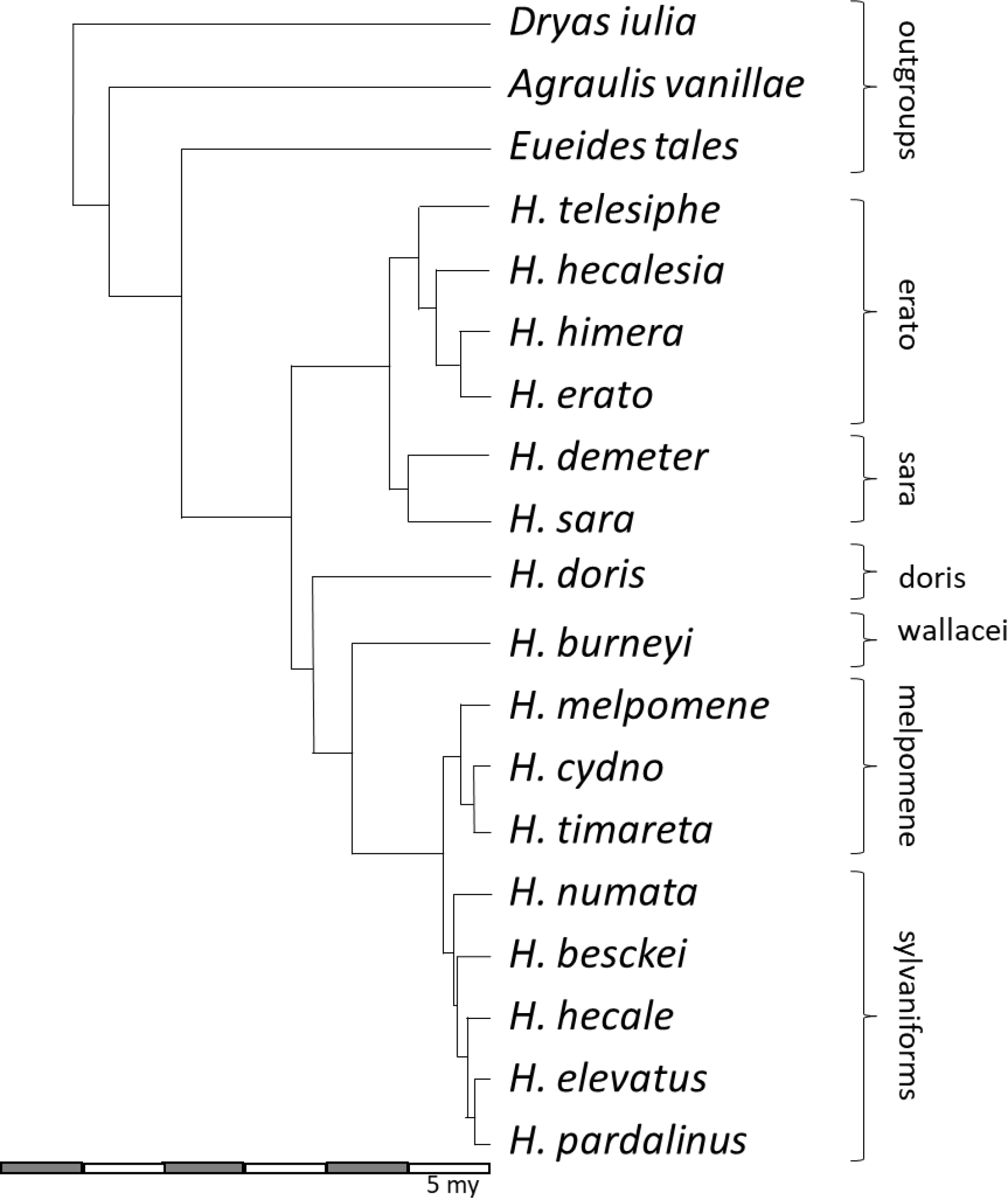
Phylogeny of the taxa examined, modified from Kozak et al. (2015). Subclade memberships are identified to the right of the tree.

### Novel and Known TE Families

After culling to eliminate duplicates and previously identified TEs, 93 novel DNA transposon consensus sequences, 59 novel LINE consensus sequences, 136 novel Helitron elements, and 65 novel LTR elements were identified. Among SINEs, the previously identified Metulj family was examined using a network-based approach (Levy, et al. 2017). That analysis yielded 2483 novel subfamilies, adding substantially to the Metulj diversity (~30 subfamilies) described in (Lavoie et al. 2013).

Three novel SINE families, which can be grouped as a new superfamily we are calling ZenoSINEs because of their presumed mobilization partner and other shared characteristics, were also identified. A fourth novel SINE family with similarities to R1 LINEs was also identified. All novel TE consensus sequences will be deposited in DFAM (Hubley, et al. 2016).

### Recent vs. Ancient Taxonomic Distributions

Because our interest was in determining how TEs may be influencing genomic structure in modern species, we distinguished between recent and ancient accumulation patterns. RepeatMasker hits with divergences <0.05 from their respective consensus sequences were considered ‘recent’ and >0.05 as ‘old’. Applying a mutation rate of 1.9 × 10^−9^ substitutions/site/generation and four generations/year (Martin, et al. 2016) and assuming minimal differences among species places this boundary at around 6.6 Mya, allowing us to focus on accumulation patterns in the melpomene-sylvaniformes and erato-sara clades as well as the terminal branches leading to *H. doris*, *H. burneyi*, and the three outgroup taxa (Figure 1). For each TE (separated by names, class, or family, depending on the level of analysis), total base coverage was calculated and divided by the total genome size to give a relative proportion (Figure 2). The figure illustrates the distinct shift from SINE dominance in ancestral accumulation patterns toward RC, LINE, and DNA dominance in the melpomene and sylvaniform clades in addition to distinct patterns in several additional species. Unpaired t-tests comparing all members of the erato-sara clade to melpomene-sylvaniform species indicates significant differences between accumulation patterns of recent SINE, LINE, DNA transposon, and rolling circle transposons insertions by class, p < 0.0001, p = 0.0229, p = 0.0078, p = 0.0008, respectively.

**Figure 2.**
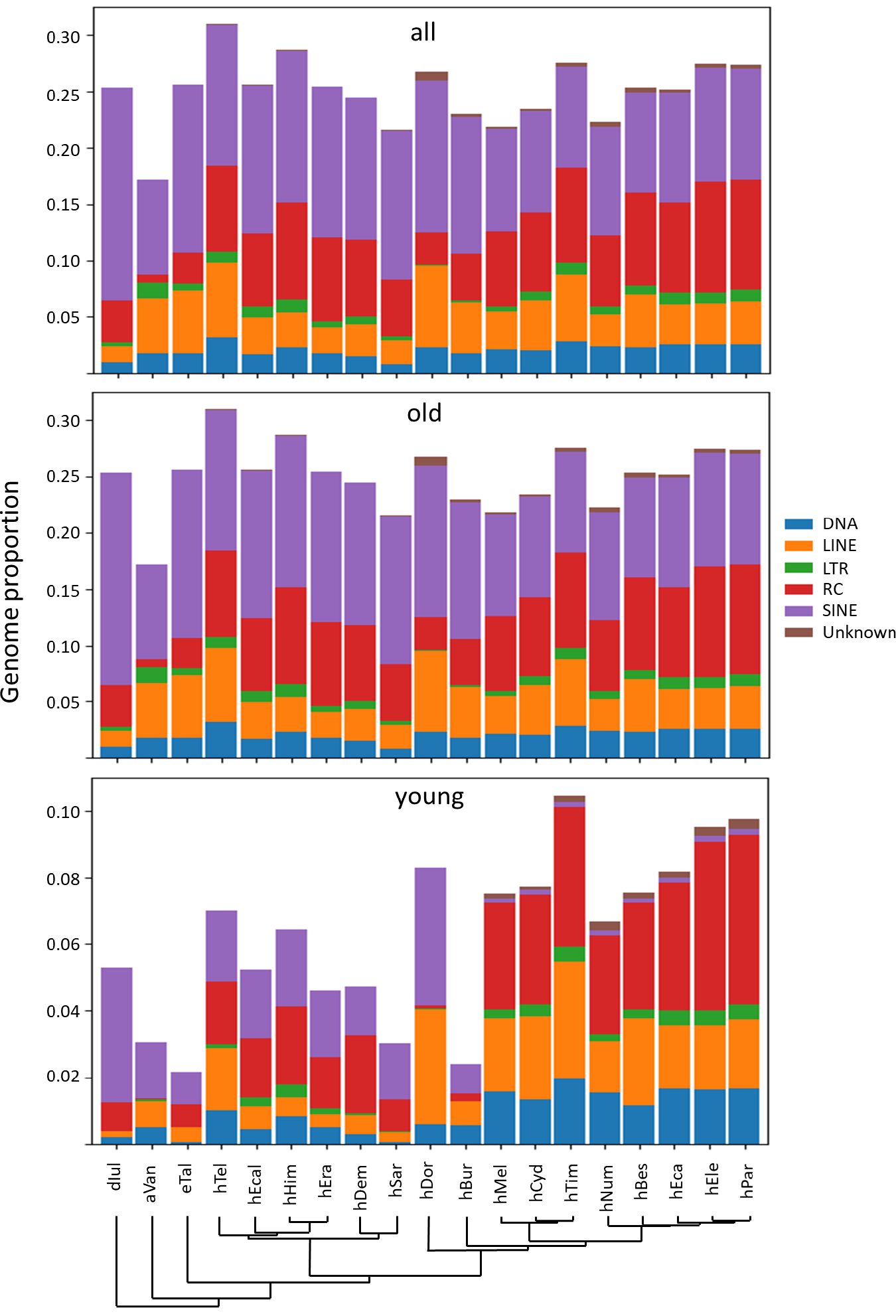
Stacked bar plots of TE proportions categorized as ‘old’ and ‘young’ in each species examined. The combined plot at the top represents ‘all’ data. Species and their phylogenetic relationships (Figure 1) are depicted on the X-axis. Values on the Y-axis are genome proportions calculated as described in the text. Abbreviations are as described in Supplemental Table 1. Briefly, the first letter indicates genus, and the following three (or four) letters, except in the cases of *H. hecale* and *H. hecalesia*, indicate species as listed in Figure 1.

Examining TE accumulation at a finer scale reveals additional patterns. For example, while recently accumulating rolling circle transposons (Helitrons) contribute to all genomes, those contributions vary substantially (Figure 3), ranging from almost no Helitron content in *Agraulis vanillae* and *H. doris* to near complete dominance in all members of the melpomene and sylvaniform clades. Further, there are clear differences in which subfamilies of Helitron have mobilized (Supplemental Figures 1 and 2). Not surprisingly, the Helitron-like elements first described by Lavoie et al. and discovered in the *H. melpomene* genome are prevalent in the melpomene and sylvaniform clades, particularly *H. elevatus* and *H. pardalinus*. Different subfamilies have recently colonized species in the erato, sara, and doris clades and to a lesser extent.

**Figure 3.**
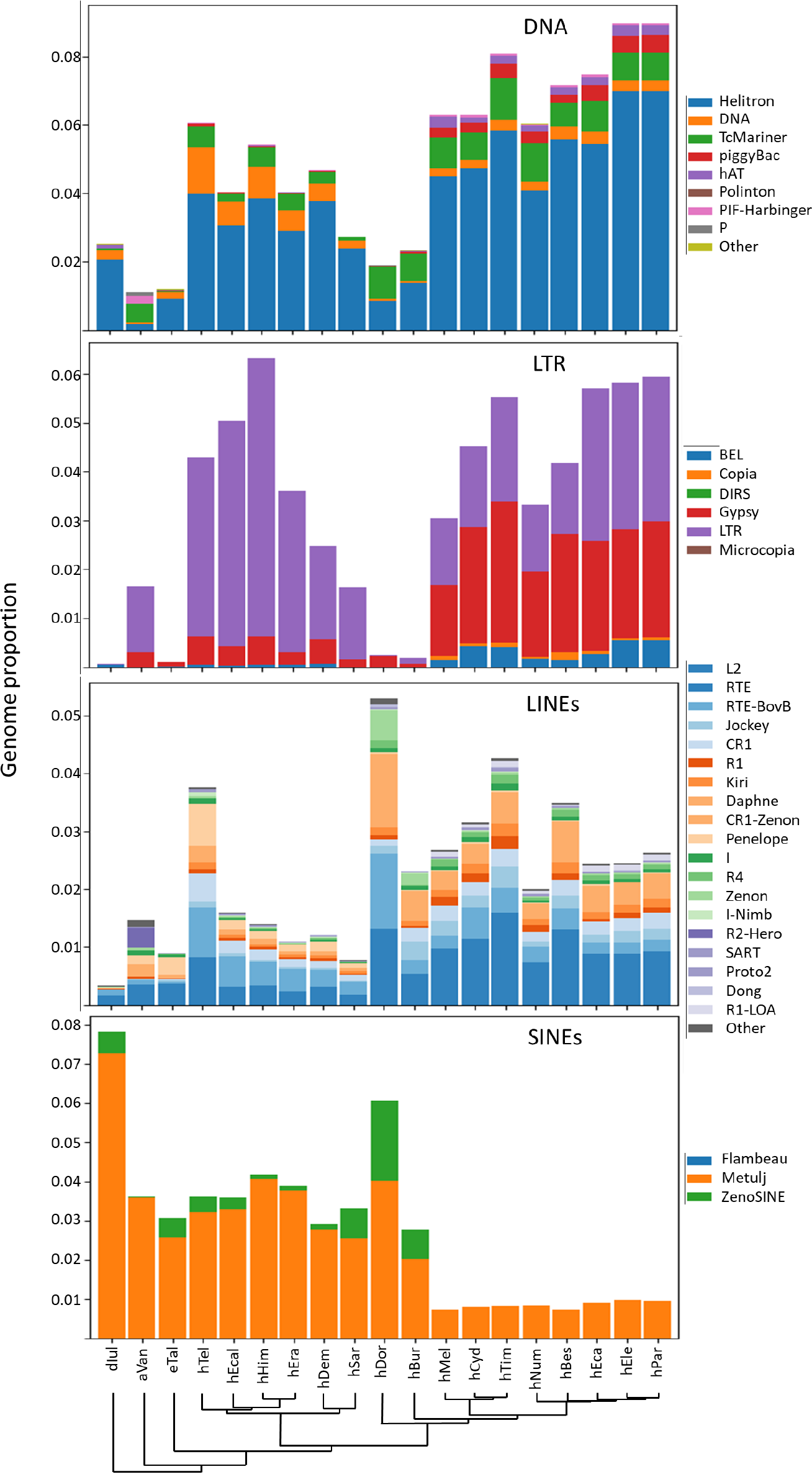
Recent contributions to genome content from each of the four TE classes examined. Axes and abbreviations are as described in Figure 2. Rolling circle (RC) transposons, (Helitrons) are depicted as part of the DNA transposon plot.

Similarly, many DNA transposons have had substantially more recent success in mobilizing within the doris, melpomene, and sylvaniform clades, again with distinct families being more prevalent, depending on the lineage (Figure 3). The most obvious difference with regard to DNA transposons lies in the increased prevalence of PIF-Harbinger, piggyBac, hAT, and most TcMariner superfamily transposons in certain clades (Figure 3, Supplemental Figures 1 and 3), especially melpomene and sylvaniform. TcMariner elements also appear to be the only DNA transposons to have managed any success in the *H. burneyi* and *H. doris* genomes while *Eueides tales* and *H. sara* seem to have avoided any substantial DNA transposon accumulation in the recent past.

Recent LTR retrotransposon accumulation patterns exhibit similar diversity (Figure 3, Supplemental Figures 1 and 4). Despite the fact there is not a significant difference in overall accumulations between members of the combined erato-sara clade and species in the combined melpomene-sylvaniform clade (unpaired t-test, p = 0.2804), there is a distinct bias toward Gypsy retrotransposons and a subset of generic LTRs in the melpomene and sylvaniform clades while a subset of LTR retrotransposons are preferred in species of the erato and sara clades as well as in *A. vanillae*. As with Helitrons and DNA transposons, the identities of the LTR retrotransposons that have expanded in each group are distinct (Supplemental Figure 4).

**Figure 4.**
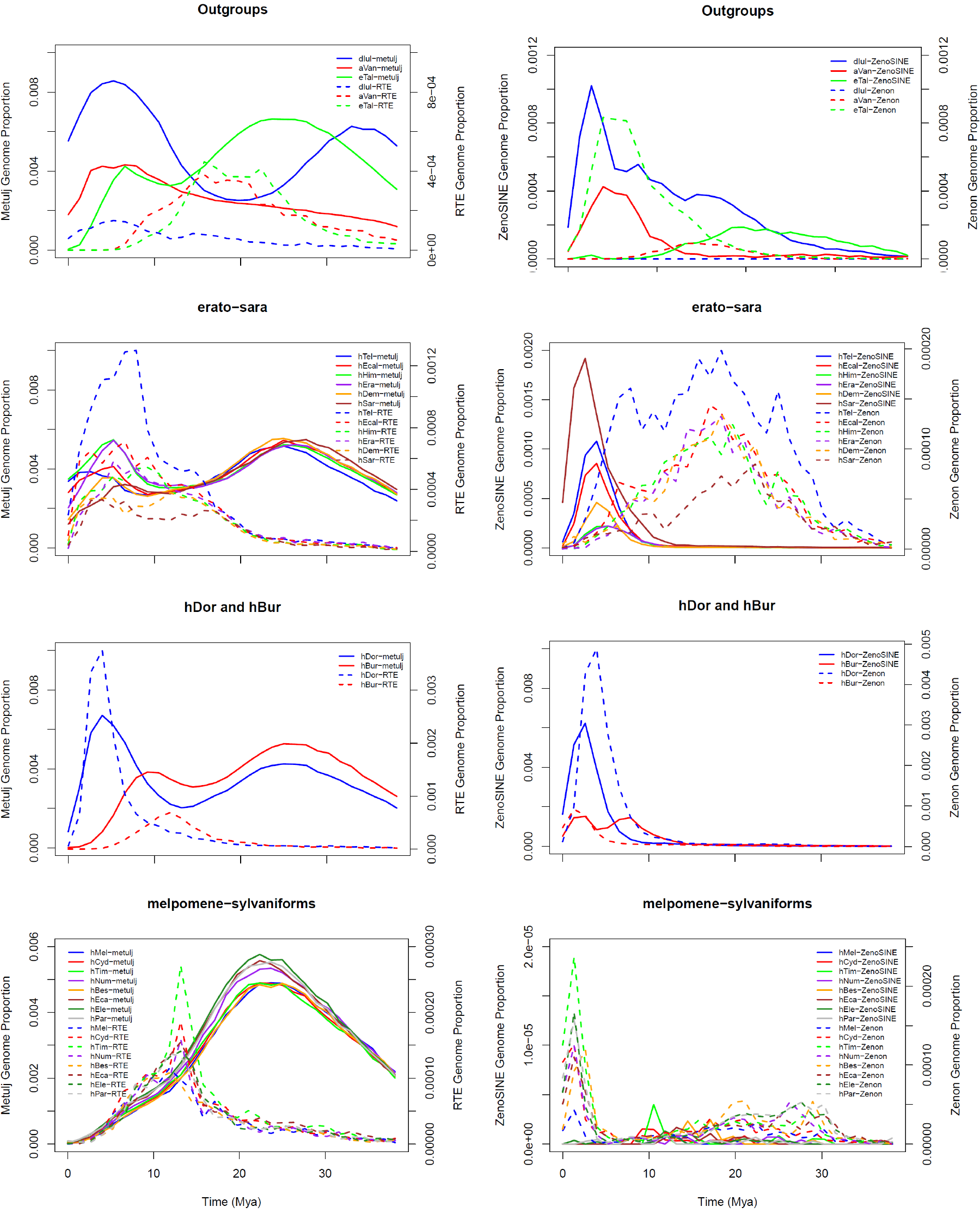
TE landscape plots for Metulj-RTE partners (left column) and ZenoSINE-Zenon partners (right column) in the four species divisions analyzed. The X-axis depicts the estimated time of accumulation of the TE using the mutation rate described in the text. Y-axes depict genome proportions for the SINE- (left axes) and LINE-derived (right axes) DNA in each genome.

Recent accumulation by LINEs is diverse but most prominent in *H. doris*, with CR1, Zenon and RTE elements dominating other LINEs (Figure 3, Supplemental Figure 5). Clades to the left in Figure 3 have generally experienced much lower levels of recent non-LTR retrotransposon accumulation. In these clades, though, a variety of short, non-autonomous Penelope elements are much more prominent, especially in *H. telesiphe*. R2-Hero elements make up a large relative proportion of LINE-occupied space in *A. vanillae*. As with the previous classes, LINE identities are highly lineage-specific (Supplemental Figures 1 and 5).

In many genomes, SINEs are the most prevalent TE component by genome proportion. As is apparent in Figure 2 and Supplemental Figure 6 this is also the case for several heliconiines. The Metulj family make the most significant recent contributions in clades other than melpomene and sylvaniform. ZenoSINEs are present only in those same clades. *H. doris* is an exception, with nearly as much accumulation from ZenoSINEs as from Metulj. Indeed, the distribution of ZenoSINEs is a puzzle. In addition to their presence in *H. doris*, and to a lesser extent *H. burneyi*, they are found primarily in *E. tales* and members of the erato and sara clades. ZenoSINEs are essentially absent from members of the melpomene and sylvaniform clades (Table 1). We examined the raw RepeatMasker output from each genome for the presence of any ZenoSINE elements greater than 100 bp in length. Counts ranged from 5-21 in the melpomene and sylvaniform clades. Sixty-two were found in *A. vanillae*, and only 12 were identifiable in *Dryas iulia*. Examination of the extracted hits on a clade by clade basis reveals that relatively few are likely to be genuine ZenoSINE elements. All of the hits from *A. vanillae* and members of the melpomene and sylvaniform clades were about half the size of the average ZenoSINE consensus, truncated at the 5’ end. For hits in the *D. iulia* genome, the hits were also short but the truncation occurred at the 3’ end. We suggest that the vast majority, if not all, such low-copy-number hits in Table 2 follow are similarly false positives.

**Table 1.** Total numbers of SINE insertions >100 bp present from each family described in the 19 genomes examined. Color coding indicates relative counts, darker green depicts higher numbers in each category.

| Taxon | Counts |  |  |  |  |  |
| --- | --- | --- | --- | --- | --- | --- |
|  | Brushfoot | Flambeau | Fritillar | Julian | Metulj | ZenoSINE |
| dlul | 385 | 134 | 10 | 16505 | 555536 | 16907 |
| aVan | 13 | 3 | 1248 | 4 | 172584 | 80 |
| eTal | 21 | 0 | 7 | 12 | 429689 | 11618 |
| hTel | 7 | 8 | 0 | 2 | 301411 | 6405 |
| hEcal | 2 | 4 | 1 | 0 | 261271 | 4172 |
| hHim | 6 | 8 | 0 | 0 | 280969 | 1492 |
| hEra | 0 | 0 | 0 | 0 | 266446 | 1440 |
| hDem | 4 | 0 | 0 | 0 | 248026 | 2012 |
| hSar | 0 | 1 | 0 | 0 | 231573 | 9139 |
| hDor | 14 | 0 | 0 | 0 | 250770 | 30999 |
| hBur | 15 | 0 | 0 | 1 | 243679 | 11912 |
| hMel | 2 | 0 | 0 | 0 | 147575 | 7 |
| hCyd | 5 | 0 | 0 | 0 | 172064 | 18 |
| hTim | 3 | 0 | 0 | 0 | 135749 | 15 |
| hNum | 4 | 0 | 0 | 0 | 200506 | 14 |
| hBes | 5 | 0 | 0 | 0 | 160502 | 25 |
| hEca | 6 | 1 | 0 | 0 | 204966 | 28 |
| hEle | 10 | 1 | 0 | 0 | 266629 | 32 |
| hPar | 6 | 0 | 0 | 0 | 232673 | 21 |

**Table 2.** TE origination rate calculations for relevant terminal and internal branches on the heliconiine tree (Figure 1). Color coding indicates relative counts and rates, darker green depicts higher numbers in each category.

| Branch | Branch Time | Threshold score | Branch-specific TEs | TE origination Rate | DNA | RC | LTR | LINE | SINE | Space contribution (Mb) |
| --- | --- | --- | --- | --- | --- | --- | --- | --- | --- | --- |
| *D. iulia* | 26.2 mya - present | 1 | 136 | 5.19 | 6 | 7 | 0 | 7 | 118 | 85.2 |
| *A. vanillae* | 23.8 mya - present | 1 | 58 | 2.44 | 7 | 2 | 1 | 22 | 29 | 35.7 |
| *E. tales* | 18.4 mya - present | 1 | 58 | 3.15 | 3 | 3 | 0 | 15 | 41 | 36.9 |
| *Heliconius* ancestral branch | 18.4 mya- 11.1 mya | 14 | 2 | 0.27 | 0 | 1 | 0 | 1 | 0 | not examined |
| erato-sara ancestral branch | 11.1 mya - 5.8 mya | 5.5 | 102 | 19.32 | 2 | 7 | 2 | 3 | 88 | 23.9 |
| erato ancestral branch | 5.8 mya - 4.7 mya | 3.5 | 9 | 1.55 | 2 | 2 | 2 | 0 | 3 | not examined |
| *H. telesiphe* | 4.7 mya - present | 1 | 3 | 0.64 | 1 | 0 | 0 | 0 | 2 | not examined |
| *H. demeter* | 5.0 mya - present | 1 | 1 | 0.20 | 0 | 0 | 0 | 0 | 1 | not examined |
| *H. sara* | 5.0 mya - present | 1 | 4 | 0.80 | 0 | 1 | 0 | 0 | 3 | not examined |
| *H. doris* | 11.1 mya - present | 1 | 104 | 9.37 | 0 | 2 | 0 | 15 | 91 | 22.3 |
| *H. burneyi* | 6.6 mya - present | 1 | 15 | 2.26 | 3 | 0 | 0 | 6 | 8 | not examined |
| melpomene-sylvaniform ancestral branch | 6.6 mya - 2.8 mya | 7.5 | 130 | 34.67 | 31 | 20 | 13 | 65 | 1 | 23.4 |

Besides ZenoSINEs, four additional new families were identified. Two of these, Flambeau, and Julian SINEs are restricted to *D. iulia*. Brushfoot is also restricted to *D. iulia* within heliconiines but has some resemblance to a possible cousin in the genome of the pierid butterfly, *Leptidea sinapsis*. Fritillar SINEs are restricted to the *A. vanillae* genome. With the exception of Julian, all are present at relatively low numbers (Table 1). Further, our analysis of the rates of evolution of new TE lineages suggests that the erato-sara common ancestor, *H. doris*, and the outgroups were hotbeds of new SINE subfamily emergence (Table 2), each associated with dozens of new subfamilies, while the melpomene and sara clades are host to a single novel subfamily.

### SINE/LINE Partnerships

The 3’ ends of SINEs are often very similar to their LINE partner (Ohshima and Okada 2005). Previous efforts by Lavoie et al. (2013)were unsuccessful in determining the LINE partner for Metulj, but based on our more complete analysis of the TE content of all 19 genomes, we now suspect that it is mobilized by an RTE family LINE (Supplemental Figure 7A). ZenoSINE, Fritillar, and Flambeau show similarity between their tails and the tail of LINEs from the Zenon family (Supplemental Figure 7B), suggesting a similar relationship. Flambeau exhibits 3’ similarity with R1 LINEs (data not shown).

The SINE tail may influence the success of the SINE in hijacking the LINE enzymatic machinery at the ribosome (Dewannieux and Heidmann 2005). Our investigations into the evolution of the 3’ tail revealed informative patterns (Supplemental Figure 8). In most *Heliconius*, young Metulj show a distinct bias toward A and T over G and C and A:T ratios are biased slightly toward T in young insertions, a signal not observed in older elements. The A prevelance over C and G is slightly higher in members of the erato and sara clades and distinctly higher in *D. iulia*, *A. vanillae*, and *H. doris*.

Because of the apparent partnership that has evolved between these SINEs and their partner LINEs, one might expect similar recent accumulation profiles. However, no relationship between the accumulation patterns is easily resolvable (Figure 4). Indeed, while there does appear to be some mirroring in *H. doris*, *H. burneyi*, and possibly in the erato and sara clades, the accumulation patterns observed in melpomene and sylvaniform are essentially opposite. A similar lack of correspondence in landscapes is apparent for ZenoSINE and Zenon LINEs. Examining correllations between recently accumulated SINEs and LINEs also reveals no discernable pattern (Supplemental Figure 9). While the expected high correspondence between ZenoSINE and Zenon LINEs is observed, so are high correlations with Dong and RTE-BovB. Further, the expected correlation between RTE-type LINEs and Metulj is not observed.

### SINE Birth and Death

Four of the novel SINEs likely originated recently within the Heliconiini. A BLAST search of all taxa excluding *Heliconius* in the NCBI WGS database using ZenoSINE consensus sequences yields only severely truncated and low similarity hits in the genomes of other lineages. Analysis of Fritillar suggested that it is restricted to *A. vanillae*, strongly suggesting that it originated in that lineage. The BLAST search produced 12 high similarity, partial hits to the fellow nymphalid butterfly *Vanessa tameamea* (the Kamehameha butterfly, GCA_002938995.1). The hits are limited to the 5’ (likely tRNA-derived) half of the SINE suggesting that these are merely hits to a similar precursor tRNA in that genome.

ZenoSINE subfamilies are similarly restricted to a subset of heliconiine lineages (Figures 3 and 4), suggesting an origin near the base of the heliconiine clade. Our BLAST search yielded hits only to *H. aoede (*GCA_900068225.1), which is sister to the doris-wallacei-melpomene-sylvaniform assemblage. Questions that will be addressed below exist on how the current distribution came to be.

Metulj are present in all species examined, suggesting that their origin is more ancient but at least prior to the diversification of the Heliconiini. A BLAST search of the NCBI WGS database yields hits only in heliconiine genomes thus far deposited with NCBI. Similar results are obtained by a broader search of all insect nucleotides in the database. Thus, while a specific point of origin cannot be identified, we suggest that Metulj originated with the clade or shortly thereafter. The lack of any substantial recent accumulation in members of the melpomene and sylvaniform clades (Figure 3) strongly suggest that Metulj is dead or dying in those lineages.

### TE origination rates

Table 2 details the rates at which various branches in the phylogeny gained novel TEs. In agreement with much of the data presented above, the erato and sara clades along with *H. doris* and the three outgroups have been home to intensive SINE diversification while the melpomene and sylvaniform clades have played host to origination events for most other categories. The highest rates of TE origination appear to center on the ancestors of each of the two major subclades and in *H. doris*.

Using this information, we determined amounts of lineage-specific TE-derived DNA contributions along selected branches of the tree (Supplemental File 1). Substantial contributions to genome diversity were observed. For example, at least 15% (~85 Mb) of the *D. iulia* genome is uniquely TE-derived when compared to any other species analyzed with most of that content (~11%, ~62 Mb) derived from lineage-specific SINEs. Around 5.5% (22 Mb) of the *H. doris* genome is unique to that lineage. Clade-specific TE contributions to the erato-sara and melpomene-sylvaniform clades are similar, averaging 5.9% (~24 Mb) and 6.9% (~23 Mb), respectively. Not surprisingly given the observations above, those contributions are quite distinct, with SINEs making up the majority of novel DNA (~15 Mb) in the erato-sara clade and DNA transposons comprising the majority (~18 Mb) in members of melpomene and sylvaniform.

### Genome size correlations

Recently, Talla et al. (2017) found that genome size in wood-white butterflies (Leptidea) correlated strongly with TE accumulation. To determine if the same phenomenon was observable in heliconiines, we followed Talla et al. (2017) and calculated a linear model of genome size as a function of TE length, and found no significant correlation (p=0.11). However, we did find a marginally significant correlation of genome size with TE count (p=0.0165). We repeated the analysis accounting for phylogenetic relatedness using the *pic* function in the R package ape v5.1 using a tree generated from concatenated, non-coding, fully-aligned regions to perform the phylogenetic correction (Edelman et al). Our results were consistent, though for both comparisons relatedness did account for some of the variation (TE length p=0.306, TE count p=0.0275). All following analyses were performed with this phylogenetic correction.

Because these species diverged very recently, we hypothesized that recent insertions may be more relevant for differences in genome size. However, this was not consistent with the data. When only considering TE insertions with divergence values less than 0.05, we found no association of genome size with either TE length (p=0.0891) or TE count (p=0.482).

To determine if any one element could be influencing genome size evolution, we next classified each insertion based on both class and family and analyzed each independently. For the full data (recent and old elements), after correcting for multiple comparisons, only I.Nimb elements were significantly associated with genome size (I.Nimb length p=5.17e^−5^, I.Nimb count p= 8.76e^−5^). However, I.Nimb elements make up only a small fraction of the genome, and the pattern appears to be driven by two outliers, *H. telesiphe* and *E. tales* (Supplemental Figure 10). For the recent elements, again a single element, Penelope, is associated with genome size (Supplemental Figure 11), but here the association is with count alone, and again it appears to be driven by the high density of Penelope in *H. telesiphe* (Penelope length p=.059, Penelope count p=1.1e^−4^).

## Discussion

TE distributions and expansion dynamics can reveal vital information about evolutionary processes. For example, taxonomic disparities in the distribution of a TE family is a sign of possible horizontal transfer among disparate lineages. The presence of high numbers of orphaned TE fragments is an indicator of high rates of non-homologous recombination that acts to remove DNA from the genome, impacting genome sizes. Thus, detailed examinations of TE content is an important step in understanding how genomes evolve.

This work is the first large-scale, comprehensive analysis of TE dynamics in a coherent clade and reveals substantial information on how heliconiine genomes diversify. Our analysis of recent accumulation patterns reveals that clear taxonomic differences have evolved with regard to the relative success of TE families across the clade.

The most obvious differences are apparent shifts in which TEs succeed in proliferating in each clade. A basal divergence in TE accumulation has evolved in *Heliconius*, with members of the melpomene and silvaniform clades showing a bias for rolling circle transposons, DNA transposons and LINEs. Meanwhile, their cousins in the erato and sara clades have been host to substantial recent SINE accumulations. Two other *Heliconius* species examined appear to have undertaken divergent strategies. *H. doris* seems to split the difference between the ‘right’ and ‘left’ clades in Figure 3 in allowing substantial accumulation from both SINEs and LINEs in the recent past while *H. burneyi* has restricted the proliferation of nearly all TEs without regard to class membership.

These observations suggest that there are substantial differences in the ways that each species deals with genomic stress caused by TE mobilization and that TE defense strategies diverge rapidly in each lineage. This is consistent with the model of piRNA clusters acting as TE ‘traps’ in which, upon an element’s insertion into a cluster, a piRNA-based defense against that element is mounted (Lu and Clark 2010). As *Heliconius* butterflies diversified, different TEs would be expected to have fallen into piRNA traps evolving in each lineage, leading to different levels of response. This would yield clade-specific patterns similar to those observed here. With the detailed descriptions we have provided, this is a hypothesis that could eventually be tested.

SINEs are often the most numerous TEs in eukaryotes. For example, while LINEs outstrip SINEs in the human genome by mass, the number of SINE insertions in our genomes surpasses LINEs by an order of magnitude (Lander, et al. 2001). With such high copy numbers, SINEs are responsible for significant structural changes and therefore deserve special attention (Wang and Kirkness 2005). SINEs are also relatively short-lived residents of many genomes, often showing higher lineage-specificity when compared to their LINE partners. This pattern is observed in the present study as we can identify all three phases of a SINE life cycle, birth, expansion, and (potentially) death.

Examination of Metulj elements suggest an interesting history. Their unambiguous presence in all species makes it clear that they evolved in the common ancestor of Heliconiini. However, their recent expansion is restricted to only a subset of the taxa examined. This suggests lineage-specific mechanisms acting to either silence this family either through active mechanisms or via self-downregulation or through massive increases in SINE mobilization. Depicting the data as TE landscapes suggests a combination of these mechanisms (Figure 4). Applying the neutral substitution rate of Martin et al. (2016) to divergence values, one can see that all members of *Heliconius* and *E. tales* experienced a peak of Metulj activity ~25 mya. This timing corresponds well with the time that a common ancestor of these species existed (Figure 1). After the initial *Heliconius* divergence, all species exhibit a decline in TE accumulation as one moves toward the present, but this is followed by resurgences in all lineages with the exception of melpomene and silvaniforms. Indeed, the lack of variability in recent Metulj content (Figure 3) suggests a rapid cessation of activity in the common ancestor of these clades.

The reason for the death of Metulj in the latter clades is unclear, as is the cause of the resurgence in other species. Why any SINE goes extinct is unknown and could be influenced by multiple factors including genomic defenses, the partner LINE, mutations in the SINEs themselves, and population genetic processes. The evolution of new subfamilies requires mobilization of the elements. Thus, the lack of any new subfamilies that are unique to this clade suggests a simple cessation of retrotransposition. If we are correct in our conclusion that RTE LINEs are responsible for Metulj mobilization, some clues may be found by examining those elements. One potential explanation is to view the SINE-LINE relationship not as a partnership but as a competition for the enzymatic machinery produced by LINEs. If the SINEs are particularly effective at hijacking that machinery, it may be possible for them to suppress LINE mobilization to some extent, even to the eventual demise of the LINE partner, as was recently hypothesized in sigmodontine rodents (Yang et al. pers. comm.). Our analysis of Metulj tails suggests that the ancestral tail of Metulj SINEs was A-rich and that a switch toward tails containing more T residues may be involved in the success of this SINE in the erato and sara clades. This hypothesis does not, however, hold true for *D. iulia*, *A. vanillae*, or *H. doris*, which have all experienced high rates of recent Metulj accumulation but exhibit a bias toward A nucleotides in their tails. These results suggest that the reasons for the differential success in heliconiine genomes may be many, and complex.

Not surprisingly, the outgroup species, with their deeper divergences, exhibit their own unique patterns. *D. iulia*, with the highest proportion of Metulj in its genome, experienced a recent surge in accumulation that outpaced any other heliconiine examined. *E. tales* mirrors the erato and sara clades while *A. vanillae* appears to have experienced a gradual increase in accumulation very recently.

Previous analyses (Lavoie, et al. 2013) suggested that larger TEs were removed via non-homologous recombination. This hypothesis is not refuted by our data. Examination of the TE landscape plots described above suggests that, unlike the pattern observed in mammalian genomes, where TEs remain as molecular fossils over large swaths of evolutionary time (Lander, et al. 2001; Waterston, et al. 2002), there is substantial turnover of TEs in these butterfly genomes. For example, when examining the temporal accumulation landscapes of Metulj, a SINE that averages well under 300 bp, we can readily see evidence of ancient accumulation (Figure 4). The LINE TE classes exhibit much less clear signatures: we rarely see ancient peaks in accumulation plots (Supplemental Figure 12). This suggests that these genomes can rapidly diverge over evolutionary time once reproductive isolation is acquired, with distinct lineages retaining little ancient TE-derived homology from larger elements across their genomes.

Assuming the phylogeny proposed by Kozak et al. (2015) and Edelman (submitted), the distribution of ZenoSINE elements is difficult to explain. The family is present at substantial numbers in *E. tales*, all members of the erato and sara clades, *H. doris*, and *H. burneyi*. Such a distribution could be explained by three scenarios. First is an ancient origin for the family in the common ancestor of the monophyletic group that includes *E. tales* and all members of *Heliconius* followed by not just a loss of activity in the melpomene and silvaniform clades but also by the removal of any previously existing insertions. The lack of any genuine ZenoSINEs (see Results) in these genomes makes the ancient origin hypothesis less likely.

Finally, it is possible that ZenoSINE evolved in only one of these lineages and this was followed by migration, either through horizontal transfer or hybridization, to the others. For example, one such scenario would be that this SINE evolved in the common ancestor of the erato and sara clades and managed to move to the other species in which it is found. Given the high tendency toward hybridization in the *Heliconius* clade overall (Mavarez, et al. 2006; Kronforst 2008; Heliconius 2012; Nadeau, et al. 2012), this seems the most plausible scenario but horizontal transfer, given that it could by a common phenomenon in insects (Peccoud, et al. 2017) cannot be ruled out.

Rates of TE origination in Heliconiini follow some expected patterns. *D. iulia*, with the longest branch on the tree has the highest fraction of branch-specific TEs (Table 2). This would be expected given a relatively constant rate of TE origination and the ancient divergence that it represents. However, examination of *Heliconius* suggests that TE origination rates are not uniform along the tree. Instead, there is a burst of TE evolution during the early stages of *Heliconius* diversification, in particular on the branch leading to the melpomene and silvaniform subclades, which spans a period ranging from ~7 – 3 mya. This corresponds well with the findings of Kozak et al. who identified a rapid increase in species diversification during the same period (Kozak, et al. 2015). Those authors proposed that environmental perturbation allowed for the invasion of new niches. This also corresponds with the periods of extensive cross-lineage hybridization found by Edelman et al. Collectively, this suggests that TEs may have been shuffled between lineages during this time. Such mixing could lead to “mismatching” in TE content vs. TE defense machinery and subsequently permitted the extensive accumulation of different TEs in different lineages. While we do not yet have data to support such a scenario, similar mismatches have been shown to play a role in *Drosophila* reproductive isolation (Petrov, et al. 1995).

TEs have been shown to respond to environmental stressors, thereby leading to substantial genomic instability (Rey, et al. 2016). Such instability has the potential, in turn, to provide novel genotypes and phenotypes upon which selection can act, either through direct changes to coding regions (Clark, et al. 2006) or through perturbations of gene regulatory pathways (Chuong, et al. 2016, 2017; Trizzino, et al. 2017). We suggest that the geologic and climatic upheaval described for this period (Gregory-Wodzicki 2000; Hoorn, et al. 2010; Jaramillo, et al. 2010; Rull 2011; Blandin and Purser 2013), may have set this cascade into motion in Heliconiini. Indeed, one recent study found that regulatory elements that differed between the sister species *H. erato* and *H. himera* were enriched for LINE content (Lewis and Reed 2018), suggesting an impact by LINEs on regulatory innovation.

Regardless, the observations presented here make it clear that differential TE activity and accumulation can act as a driver of rapid genomic divergence. Similar analyses of multiple taxa have been performed for other groups including squamates and birds (Kapusta and Suh 2017; Pasquesi, et al. 2018). In those studies, especially the squamates, similar shifts in TE content and accumulation were observed. However, those analyses examined much deeper divergences than the ones examined here. Thus, one might expect to observe more drastic changes because of the longer evolutionary time spans. In examining much more closely related lineages, we demonstrate that even over relatively short periods, the TE landscapes in members of a single genus can diverge rapidly due to differential TE dynamics. Lineages whose common ancestor harbored a single complement of TEs now play host to very distinctive complements of recently active TEs with patterns that resemble genomic fingerprints. Even in the case of LTR accumulation, where no significant difference exists with regard to overall accumulation amounts, the identities of the elements that have accumulated are quite distinct. Such distinctions are true of all classes. This is exemplified by our observation that on average ~23 Mb (5.3-9.2%, depending on genome size) of the genomes of the melpomene and sylvaniforms subclades harbor TE-derived DNA that would not be found in members of erato and sara. In *D. iulia*, a full 15% (85.2 Mb) of the genome is uniquely TE-derived in that lineage when compared to any other species we examined. The data make it clear that novel TE families, such as ZenoSINE and Julian, can arise and replicate rapidly to occupy substantial genome fractions in isolated lineages. Furthermore, because these genomes tend to actively remove longer TEs, the ancestral fractions of each genome will change rapidly as different portions are removed in each lineage.

This purely structural component of genome evolution, when combined with the functional impacts of TEs as they contribute new open reading frames, regulatory sites, and small RNAs add support to the contention that TEs are major drivers of genome evolution and deserve significant attention when determining the forces that lead to the taxonomic and phenotypic diversity around us.

These results also suggest powerful ways to move forward in understanding the forces that act to regulate TE activity. Here, we provide what amounts to a ‘natural history’ of TEs content in the genomes of 19 relatively closely related species. Researchers interested in how small RNAs and their protein partners act to suppress the damage of TEs now have a detailed starting point from which to begin detailed studies.

## Materials and Methods

### TE discovery and Classification

De novo TE discovery was implemented using a combination of RepeatMasker (Smit, et al. 2013-2015), RepeatModeler (Smit and Hubley 2008-2010), and manual annotation as described in Platt et al. (2016) with some modification. Briefly, each genome assembly was sorted by scaffold length and the top ~200 Mb were used as the base for our analysis. Each genome fragment was then subjected to a RepeatModeler analysis and a de novo repeat library was generated. Each genome fragment was then masked using its de novo library. RepeatMasker output was processed using a custom Perl script to calculate K2P distances for each insertion.

Because our primary interest is in lineage-specific insertion patterns, we sorted insertions by K2P distance from their respective consensus sequence and selected only insertions that were likely to be recent. K2P distance cutoff values were determined using information from the phylogeny of Kozak et al. (2015). For example, several subgroups are evident from the phylogeny in Figure 1. Three species form a relatively deeply diverged set of outgroup taxa, *A. vanillae*, *D. iulia*, and *E. tales*. Because of the longer branch lengths, these species are likely to harbor older but still lineage-specific insertions compared to species in the more recently diversified clades. We therefore examined any insertions with divergences <0.2 in the outgroups. Similarly, we used reduced cutoffs for members of the other three groups (i.e. divergences <0.1 for members of the doris and wallacei clades and <0.05 for members of the erato, sara, melpomene, and sylvaniform clades).

Manual validation of putative repeats discovered by RepeatModeler was performed as described in Platt et al. (2016) by using them as queries against a combined ‘pseudogenome’ consisting of a concatenation of each 200 Mb fragment draft with BLASTn v2.2.27 (Altschul, et al. 1990). Repeats with fewer than ten hits were discarded from downstream analyses. For all remaining queries, the top hits (up to 40) were extracted with at least 500 bases of flanking sequence and aligned with the query using MUSCLE v3.8.1551 (Edgar 2004). Majority rule consensus sequences were generated in BioEdit v7.2.5 (Hall 1999) and manually edited to confirm gaps and ambiguous bases. 5’ and 3’ ends were examined for single copy DNA, indicating element boundaries. If no single copy DNA was identifiable, the new consensus was subjected to new iterations until boundaries were detected. After each round, new consensus sequences were subjected to a consolidation check using cd-hit-est (Li and Godzik 2006) to identify consensus sequences that could be combined. Criteria for collapsing two or more consensus sequences were 90% identity over at least 90% of their total length.

Broad categories of TE (i.e. DNA transposons, rolling circle transposons (RC), LINEs, SINEs, LTR elements, and unknown) were determined using a combination of BLAST searches of the NCBI database and CENSOR searches of Repbase (Jurka, et al. 2005; Kohany, et al. 2006). We also used structural criteria as follows: for DNA transposons, only elements with visible terminal inverted repeats were retained. For rolling circle transposons we required elements to have an identifiable ACTAG at one end. Putative novel SINEs were inspected for a repetitive tail and A and B boxes. LTR retrotransposons were required to have recognizable hallmarks such as TG, TGT or TGTT at their 5’ and the inverse at the 3’ ends. Because of the complexity of SINE evolution, putative SINEs were analyzed uniquely as described below. While sequences in the unknown category could be transposable elements, they formed only a very small fraction of the total putative TE sequence, and they could also represent segmental duplications or other non-TE species. Our interest was in the TE dynamics in these genomes, thus, these were ignored in most downstream analysis. All other categories were checked for high similarity to known TEs and to one another using a final combined run of cd-hit-est using the same criteria as previous.

### SINEs

SINE evolution is complex and identifying subfamily structure is a difficult problem, primarily due to the high number of insertions typical of a genome. Initial analysis suggested three SINE families in these genomes. The first is the previously described Metulj family. The second is a novel family that appears to be derived from the fusion of Zenon LINE 3' tails with a 5’ head of unknown origin, which we call ZenoSINE. A small subfamily distinct from the main ZenoSINE family was identified in and restricted to the *A. vanillae* genome *A. vanillae* is commonly known as the Gulf Fritillary. Thus, we dubbed this subfamily ‘Fritillar’. Finally, a third family that is derived from R1 elements is restricted to *D. iulia*. One common moniker for this species is ‘flambeau’ and we suggest the same name, Flambeau, for this family of SINEs.

Metulj SINEs were far more numerous and widespread than their ZenoSINE cousins (discussed below), and therefore represented a more difficult analytical problem. A recently developed network-based method for subfamily (aka community) detection was used to identify Metulj subfamilies (Levy, et al. 2017). Briefly, similarity networks were constructed by pairwise-aligning Metulj elements >240 nucleotides long (n = 498,141) from all 19 butterfly genomes using BLAST. Further preprocessing was performed to prevent possible biases caused by sequence length and shared poly(A/T) tails that may confound community detection. For this step, previously identified Metulj consensus sequences were aligned using MUSCLE and 5’ and 3’ overhangs were manually trimmed using Bioedit. Genomic Metulj sequences were aligned to these trimmed consensus sequences using BLAST+ to identify corresponding regions (parameters: -*strand plus -max_target_seqs 3 -num_threads 20 -word_size 4 -evalue 1e-2 -dust no -soft_masking false*). Minimum start and maximum end positions define the region for further analysis per sequence and were length-filtered for >=235 nucleotides. The 420,689 sequences retained were analyzed for subfamily detection: the sequences were pairwise aligned using BLAST (version 2.7.1+; blastn command was used with non-default parameters: -*strand plus -dust no -max_target_seqs 50 -word_size 8 -soft_masking false*). Bornholdt community detection (Reichardt and Bornholdt 2006) was applied using *gamma*=*59*. Consensus sequences were computed using MUSCLE with 30 randomly selected sequences per community (with max of 2 iterations). To further refine subfamily definitions, communities with identical consensus sequences were merged (such pairs were identified using BLAST requiring 100% identity and 95% query coverage). Consensus sequences were computed per subfamily and were used to refine the subfamily annotation, resulting in a final set of 2,493 subfamilies (Supplemental File 2). This set was further grouped into 147 clusters to simplify downstream analyses using cd-hit-est. Clustering criterion was 95% identity, comparing the entire length of the SINEs

### LINEs

Previous analyses (Lavoie, et al. 2013) suggest that longer TEs are more likely to be fragmented by non-homologous recombination. As a result, we focused on the LINE open reading frame to increase the potential for comparable data. A special effort was made to identify full- or near full-length open reading frames (ORFs) for each clade. First, we identified all known LINE elements from the *H. melpomene* genome in RepBase. These were combined with any LINEs identified in our de novo analysis after removing possible duplicates. All remaining elements were filtered, retaining any with intact ORFs of at least 2kb, starting with methionine, and with clearly identifiable start and stop codons using ‘getorf’ from the EMBOSS package (Rice, et al. 2000).

To identify subfamily structure of LINEs, phylogenetic analysis of these ORFs was accomplished by masking each genome with the resulting library and retaining any hits of 1.5kb or longer. Generally, extracted hits were aligned using MUSCLE and subjected to a neighbor-joining (NJ) analysis (described below). However, large numbers of hits impeded efficient alignment in some cases due to memory limitations. To work around this problem, we reduced the number of hits by randomly selecting smaller numbers of sequences from the pool and re-aligning until successful. In some of these cases, there was a lack of overlapping sites that impeded the NJ analysis. In these cases, we extended our filter to include hits that were at least 2kb, producing the needed overlapping regions.

Each set of aligned ORFs was subjected to NJ analysis to identify any apparent structure. NJ analyses were accomplished based on the maximum composite likelihood parameters in MEGA7 (Kumar, et al. 2016) with pairwise deletion of ambiguous positions and 500 bootstrap replicates. Trees were examined visually and clearly delineated clades with high bootstrap support were labeled as subfamilies using letter designations (Supplemental File 2 and Supplemental Figure 13). For example, examination of the RTE-4_Hmel tree yielded four subfamilies, RTE-4_Hmel_A-D (Supplemental Figure 13).

To estimate genetic distances among members of each subfamily, we used a combination of tools via a custom script that would first align the hits identified for each subfamily using MUSCLE. The script would then invoke trimal (-gt 0.6 -cons 60 -fasta) to trim the alignment (Capella-Gutierrez, et al. 2009) and use ‘cons’ from the EMBOSS package to generate a consensus sequences (-plurality 3 -identity 3). We then used MEGA7 to calculate mean divergence from the consensus, mean divergence among subfamily members, and divergence ranges (Supplemental File 3).

### Recent vs. Ancient Taxonomic Distributions

To determine taxonomic distributions for each class, family, and subfamily, we used RepeatMasker and custom python scripts to generate proportion tables as follows. RepeatMasker was used to identify insertions in each of the 19 genomes, this time using the entire genome drafts. Hits with divergences <0.05 from their respective consensus sequences were considered ‘recent’ and >0.05 as ‘old’. For each TE (separated by names, class, or family, depending on the level of analysis), total base coverage was calculated and divided by the total genome size to give a proportion.

To illustrate differences among *Heliconius spp*. In terms of TE composition, we imposed a principal components (PC) analysis on a species-by-element matrix each for DNA transposons, LRT transposons, SINE’s and LINE’s. To illustrate similarities and differences among Heliconiini, we displayed their positions based on the first two principal components. Species that are proximate in this two-dimensional space have more similar TE composition than species that are more distant. To illustrate how species differed based on their TE composition, we displayed the correlation of each individual element type (e.g. those with unique names) with the first and second PC.

### SINE/LINE partnerships

SINEs and LINEs have a host-parasite relationship with SINEs, in which SINEs will hijack the enzymatic machinery encoded by their partner LINE to mobilize (Kajikawa and Okada 2002; Roy-Engel, et al. 2002; Dewannieux, et al. 2003). Such partnerships are often defined by a shared 3’ tail (Ohshima and Okada 2005). We examined the 3’ ~100 bp of each SINE and queried the 3’ ends of all LINEs in our new TE database to determine the likely LINE partner for each.

The 3’ tails of Metulj elements exhibited substantial complexity, with a variety of structures including poly-A tracts, poly-T tracts, repeated ATTTA motifs, and repeated GATG motifs, among several others. Based on previous work, we suspected that differences in the tail may influence relative success in retrotransposition (Dewannieux and Heidmann 2005; Ohshima and Okada 2005). To investigate how tail structure evolved, we extracted 100 random full-length Metulj insertions from each taxon. Each set of extracts was aligned to representative consensus sequences. This was repeated ten times for each taxon. The 3’ ends of each alignment were degapped starting where the tail begins and the ratios of each pair of nucleotides was identified and plotted after log-transformation. This was conducted separately for ‘old’ and ‘young’ SINEs.

To determine if either Metulj or ZenoSINE accumulation patterns were correlated with any LINE elements, Pearson correlation coefficients based on proportion of each genome occupied were visualized using the “corrplot” package in R and RStudio v1.0.143.

### TE origination rates

To estimate approximate rates that lineages evolved new TE lineages, we calculated the number of branch-specific TEs using RepeatMasker output. A TE was scored as ‘present’ (score = 1) in a genome if at least 5000 bp of sequence attributable to that TE was identifiable in the genomes of terminal branches. A TE was considered ‘absent’ (score = 0) if fewer than 500 bp was identified. To score subclades, we allowed ‘possible presence’ scores of 0.5 if base counts fell between the two values. Subclade ‘presence’ sum threshold scores were subclade specific based on the number of species examined. For example, the erato subclade, with four members, had a presence sum threshold of 3.5. Branch times were obtained using the median scores for each node calculated using TimeTree (Kumar, et al. 2017). Rates of TE origination were calculated by dividing the number of branch-specific insertions by the time that the branch likely existed.

We estimated lineage-specific DNA contributions to selected branches of the tree by identifying DNA that was deposited by novel TEs that evolved on those branches. We then calculated both the genome proportions occupied by those elements and the total bp. For example, we summed the total contributions made by each of the 118 novel TEs identified in the *D. iulia* genome (Table 2). Similarly, we summed total the total bp deposited by each novel TE identified on the erato-sara common branch in each member of those clades and calculated the mean (Supplemental File 1).

### Genome size correlations

Using the annotations generated, we compiled summary statistics of transposable element content in each heliconiine genome, in terms of TE bases per base pair (TE length) and number of insertions per base pair (TE count). We obtained genome size estimates from Edelman *et al.* (submitted). Because the absolute values of these measures are several orders of magnitude apart, we Z-transformed each category by subtracting the mean and dividing by the standard deviation.

## Supporting information

Supplemental file information with download links

## Acknowledgments

The authors wish to thank the College of Arts and Sciences at Texas Tech University for funding related to this work. In addition, we would like to thank the Texas Tech HPCC (http://www.depts.ttu.edu/hpcc/) for providing computational resources necessary to complete this project. Angela Peace provided assistance with early conceptual analyses. The 20-genome *Heliconius* project was funded by a SPARC Grant from the Broad Institute of Harvard and MIT as well as startup and studentship funds from Harvard University.

