## Supplemental file information with download links for "Analysis of 19 Heliconiine Butterflies Shows Rapid TE-based Diversification and Multiple SINE Births and Deaths"

**Supplemental Files.**

**Supplemental Table 1.** Taxon and genome information for the species examined. Available at myweb.ttu.edu/daray/heliconius/Supplemental_Table_1_- _Genome_info.xlsx

**Supplemental File 1.** Lineage-specific TE families and subfamilies and genome content contributions. Available at myweb.ttu.edu/daray/heliconius/Supplemental_File_1_-_Novel_TE_contributions.xlsx

**Supplemental File 2.** Metulj1_communities.xlsx available at myweb.ttu.edu/daray/heliconius/Supplemental_File_2_-_Metulj1_communities.xlsx

**Supplemental File 3.** LINE subfamily memberships and divergences.xlsx available at myweb.ttu.edu/daray/heliconius/Supplemental_File_3_- _LINE_subfamily_memberships_and_divergences.xlsx

**Supplemental Figure 1.** Variation in TE composition among species and species groups.  (Top row) Positions of species in a two-dimensional space defined by the first two principal components extracted from a species by element matrix. Plots demonstrate that particular TE families tend to be unique to each butterfly lineage. Interestingly, *H. doris* and *H. burneyi*, members of the doris and wallacei lineages, respectively, tend to cluster with the outgroups and the erato and sara lineages rather than melpomene and sylvaniforms. This suggest an overall TE-based similarity with the former lineages rather than the latter. (Bottom row) Correlations of each of the individual element families with the first two principal components. Each dot represents a distinct, named TE family or subfamily. Plots illustrate the tendency of most of the distinct TE families/subfamilies to occupy particular subsets of Heliconiine genomes. Available at myweb.ttu.edu/daray/heliconius/Supplemental_Figure_1_-_PCAs.pdf

**Supplemental Figure 2.** Heatmap of Helitron-occupied genome proportions by Helitron name. Available online at myweb.ttu.edu/daray/heliconius/Supplemental_Figure_2_-_RCpropheatmap.pdf

**Supplemental Figure 3.** Heatmap of DNA transposon-occupied genome proportions by TE name. Available online at myweb.ttu.edu/daray/heliconius/Supplemental_Figure_3_-_DNApropheatmap.pdf

**Supplemental Figure 4.** Heatmap of LTR retrotransposon-occupied genome proportions by name. Available online at myweb.ttu.edu/daray/heliconius/Supplemental_Figure_4_-_LTRpropheatmap.pdf

**Supplemental Figure 5.** Heatmap of LINE retrotransposon-occupied genome proportions by name. Available online at myweb.ttu.edu/daray/heliconius/Supplemental_Figure_5_-_LINEpropheatmap.pdf

**Supplemental Figure 6.** Heatmap of SINE retrotransposon-occupied genome proportions by name. Available at myweb.ttu.edu/daray/heliconius/Supplemental_Figure_6_-_SINEpropheatmap.pdf

**Supplemental Figure 7.** A) Alignment of selected Metulj 3’ ends without tails, which vary among subfamilies, with selected RTE and RTE-BovB LINE tails. B) Alignment of selected ZenoSINE-like tails with selected CR1-Zenon tails. Available at myweb.ttu.edu/daray/heliconius/Supplemental_Figure_7_SINE-LINE_Tail_alignments.pdf

**Supplemental Figure 8.** Boxplots illustrating 3’ tail content of old and young Metulj SINEs. Available at myweb.ttu.edu/daray/heliconius/Supplemental_Figure_8_-_Boxplots.pdf

**Supplemental Figure 9.** Heatmap of correlations between recently accumulated Metulj and ZenoSINE SINEs across the 19 genomes analyzed. Available at myweb.ttu.edu/daray/heliconius/Supplemental_Figure_9_-_Correlation_heatmap.pdf

**Supplemental Figure 10.** Correlation plot illustrating genome size vs. I-Nimb LINE content. myweb.ttu.edu/daray/heliconius/Supplemental_Figure_10_-_I-Nimb_correlation.jpeg

**Supplemental Figure 11.** Correlation plot illustrating genome sizes vs. Penelope LINE content. Available at myweb.ttu.edu/daray/heliconius/Supplemental_Figure_11_-_Penelope_correlation.jpeg

**Supplemental Figure 12.** TE landscape plots illustrating LINE accumulation patterns. Available at myweb.ttu.edu/daray/heliconius/Supplemental_Figure_12_-_LINE_plots.pdf

**Supplemental Figure 13.** Example of subfamily memberships derived from observations of NJ trees of LINE ORFs. RTE-4_Hmel, subfamilies A-D indicated by labeled colored boxes for each subfamily. Available at myweb.ttu.edu/daray/heliconius/Supplemental_Figure_13_-_RTE-4.pdf
